# Multidrug-resistant *Rhodococcus equi* (MDR-RE): detection in East Asia confirms global spread

**DOI:** 10.1101/2025.10.21.683145

**Authors:** Jorge Val-Calvo, Yuta Kinoshita, James Gibbons, Mariela Scortti, Hidekazu Niwa, José A. Vázquez-Boland

## Abstract

Macrolide (multidrug)-resistant *Rhodococcus equi* (MDR-RE) emerged in the early 2000s and has since been disseminating across equine farms in the United States. In 2016, it was found in Ireland, presumably introduced through horse imports. Here, we report its identification in Japan, together with evidence of its presence in China. A recent isolate from Ireland suggests ongoing importation events. These findings highlight the continued international spread of MDR-RE and its potential to become globally established. Currently circulating among horses, MDR-RE is likely to arise in humans over time. Active surveillance is essential to monitor and control this emerging zoonotic antimicrobial resistance threat worldwide.

---

*Rhodococcus equi* is a soil-borne pathogenic actinomycete that causes pyogranulomatous infections in a range of mammalian hosts [1,2]. The bacterium is commonly found in the farm environment, from which it is zoonotically transmitted to humans, causing rare but severe opportunistic infections. In animals, rhodococcal disease is most frequently seen in horses, where it is a primary cause of fatal pneumonia in foals [1-4]. Recently, the emergence of a multidrug-resistant (MDR) variant of *R. equi* (MDR-RE) was reported in equines [5,6]. Most MDR-RE isolates belong to a clonal lineage named “2287” [6-9] that likely arose in the US in the late 1990s /early 2000s on horse farms practising mass antibioprophylaxis against endemic *R. equi* [10,11].

MDR-RE resulted from the co-acquisition of a macrolide resistance plasmid, pRErm46, and a chromosomal rifampicin-resistance mutation, *rpoB*^S531F^ [6]. pRErm46 carries *erm*(46), an MDR-RE-specific macrolide-lincosamide-streptogramin B (MLSB) resistance determinant, on a mobilizable transposon, Tn*RErm46* [5,6]. Via its IS*481*-family transposase, Tn*RErm46* can integrate into the *R. equi* genome, including the pVAPA virulence plasmid [6]. Some members of the MDR-RE 2287 clonal complex have lost pRErm46 but retain macrolide resistance due to genomic Tn*RErm46* copies [7,8] (see strains indicated with empty circles in **Figs. 1** and **2**). Additionally, pRErm46 harbours a class I integron (C1I) carrying resistance cassettes *aadA* (streptomycin and spectinomycin), *sul1* (sulfonamides), and *tetRA*(33) (tetracyclines) [6]. This confers high-level resistance to multiple antimicrobials, including the standard macrolide-rifampicin combination therapy for *R. equi* in foals (7,12), used for over 40 years [13-15].

**Fig. 1.**
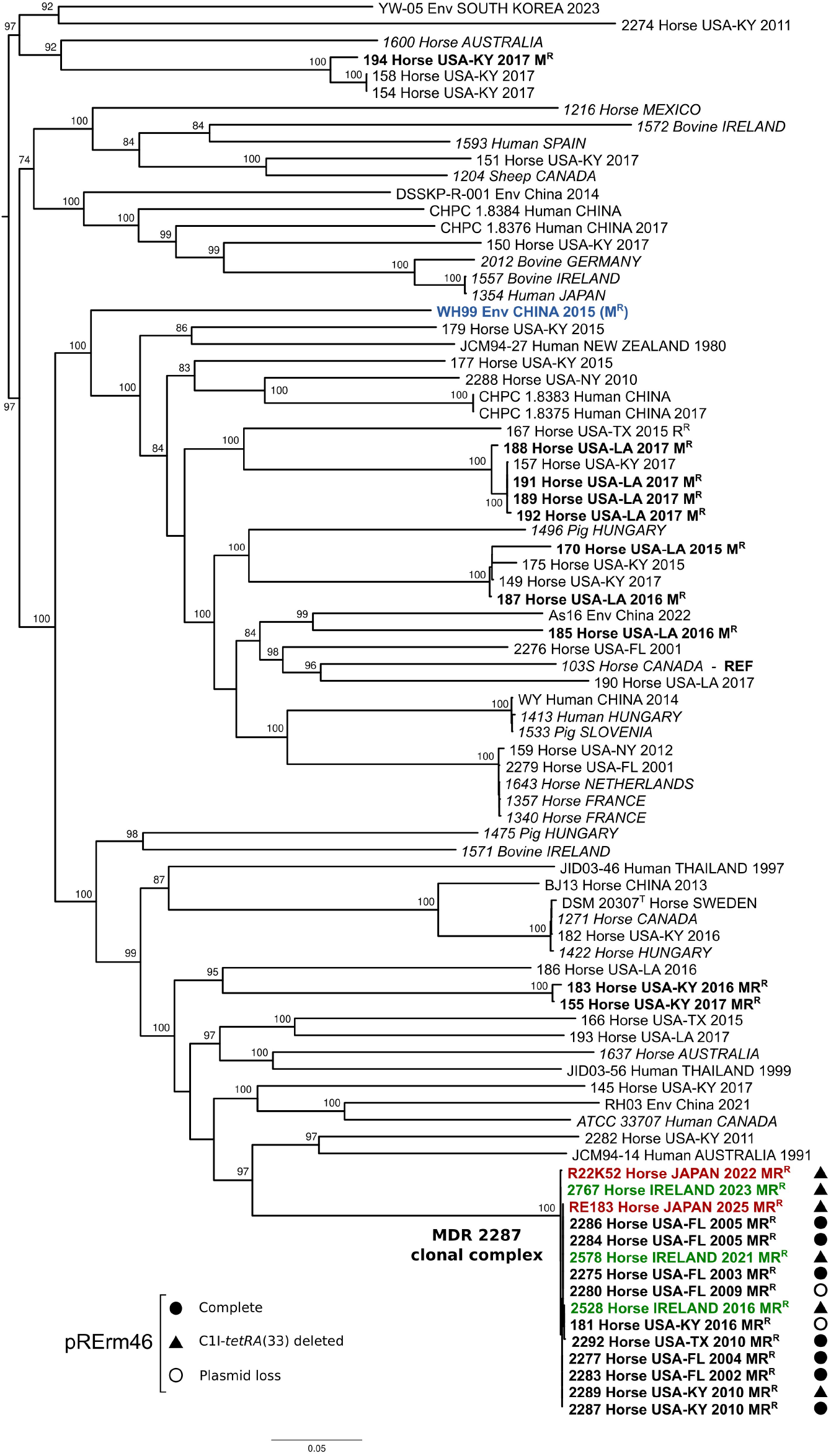
Core-SNP ML phylogeny of *Rhodococcus equi* and MDR-RE. Core genome alignment and SNP detection was performed using Parsnp v. 2.0.5. with *R. equi* strain 103S (GenBank accession no. FN563149) [24] as the reference (indicated “REF” in bold). The ML tree was inferred using IQ-TREE2 v2.4.0 (https://iqtree.github.io) with the GTR+F+ASC+R8 substitution model and rooted at the midpoint. Ultrafast bootstrap values ≥70% for 1,000 replicates are indicated. Tree drawn with FigTree (http://tree.bio.ed.ac.uk/software/figtree). A total of 84 *R. equi* genome assemblies were used, including 42 from macrolide-resistant and -susceptible equine isolates originating from the US, 21 from a previously reported *R. equi* diversity set (labels in italics) [8], and 17 from isolates from Asia or Oceania. The MDR-RE 2287 clonal complex is represented by 10 isolates from the US (in bold) [7], the three isolates from Ireland [8,9] (bold green), and the two isolates from Japan (bold red). Other macrolide-resistant isolates from different *R. equi* lineages, which have acquired the MDR plasmid pRErm46, are highlighted in bold. The WH99 soil isolate from China [18,19], where Tn*RErm46* was identified, is indicated in bold blue. Labels indicate geographic origin, year of isolation, and resistance phenotype when applicable (MR^R^, macrolide and rifampicin resistance; M^R^, macrolide-only resistance; R^R^, rifampicin-only resistance). Symbols indicate pRErm46 plasmid carriage in MR^R^ or M^R^ isolates, as described in the key (solid circles, complete plasmid; solid triangles, plasmid with ΔClI-*tetRA*(33) deletion; open circles, plasmid loss after Tn*RErm46* transposition to chromosome).

With no proven alternative therapies available for foal rhodococcosis, MDR-RE 2287 poses a serious threat to the equine industry. Around 10% of human infections involve equine-derived (pVAPA-harbouring) strains, while ≈50% are linked to porcine-derived (pVAPB-harbouring) strains (ref. [16] and unpublished data), which *in vitro* studies showed can also acquire pRErm46 [17]. Moreover, pRErm46 can conjugally transfer and persist in various related actinomycetes [17]. Thus, MDR-RE also poses a public health risk either through potential direct zoonotic transmission or as a source of resistance gene transfer for other pathogens.

## MDR-RE international surveillance

Since emerging, MDR-RE 2287 has spread across US horse-breeding farms, likely via carrier horses [7], with high prevalence in Kentucky, its presumed origin [7,10-12]. Due to concerns about international spread, an informal surveillance network was established involving 16 sentinel laboratories in 14 countries across six continents, with Edinburgh as reference centre for *R. equi* phylogenomics [8,9]. Participants prospectively monitor for *R. equi* isolates showing significant macrolide resistance (MIC ≥4 μg/ml), and retrospectively in archived isolates.

The first confirmed detection of MDR-RE 2287 outside the US was in 2016 in Ireland [8], a country with strong horse trade links to the US. Additional Irish MDR-RE 2287 isolates were recovered in 2021 and in 2023 [8,9], the latter (PAM 2767) reported in this study (**Fig. 1**). Following these, the MDR-RE surveillance initiative has now identified MDR-RE 2287 in Japan, another major horse-breeding country.

## MDR-RE 2287 in Japan

Two macrolide-rifampicin resistant *R. equi* strains, R22K52 and RE183, were submitted by Japan’s Equine Research Institute (ERI) for phylogenomic analysis. They were isolated from foal pneumonia cases in 2022 and 2025, on different farms, and were deposited in the Edinburgh collection as PAM 2822 and PAM 2831. Genomic DNA from both isolates, as well as the third Irish strain, was sequenced by Illumina (Novogen UK, 150-bp paired-end reads), and read libraries checked, trimmed and assembled, as previously described (*8*). Phred scores were 36-39, and average sequencing depth was ≥250×. Maximum Likelihood (ML) phylogenies based on conserved SNPs were used to determine the phylogenetic relationships with the *R. equi* population (**Fig. 1**) and the MDR-RE 2287 clonal complex (**Fig. 2**). Methodological details of the phylogenetic analyses as well as other bioinformatic methods can be found in the corresponding figure legends.

**Fig 2.**
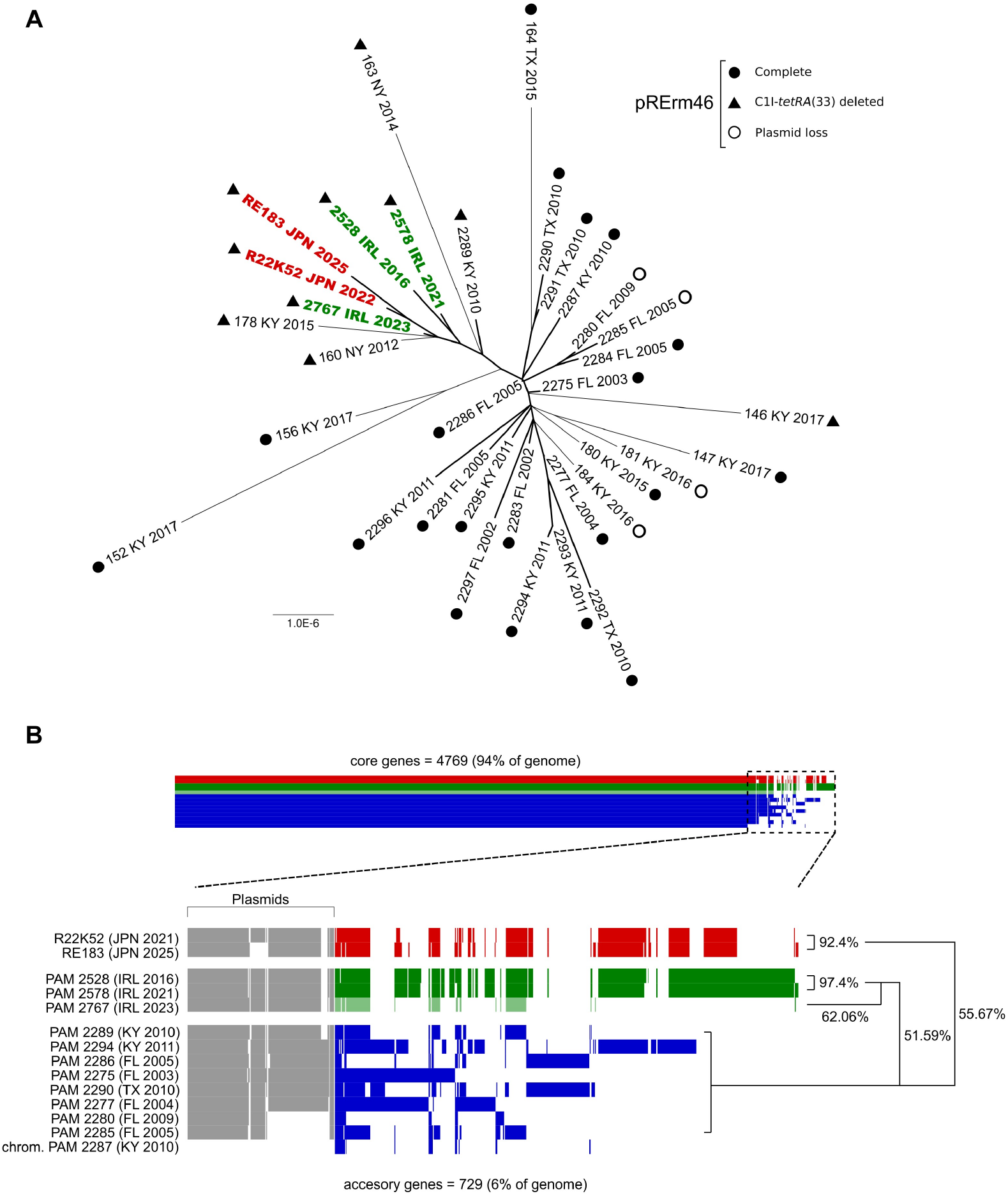
WGS analysis of MDR-RE 2287 isolates (in red, from Japan; in green, from Ireland). (**A**) *R. equi* MDR-RE 2287 ML phylogeny. Unrooted tree inferred from core SNPs called by SNIPPY (https://github.com/tseemann/snippy) using Illumina paired reads (four-digit isolate designations, thicker branches) or previously reported PacBio sequences [7] (three-digit isolate designations, thinner branches), with MDR-RE 2287 isolate PAM 2287 as the reference. PacBio assemblies with unusually high SNP counts (>50) were excluded. The tree was built using IQ-TREE2 v. 2.4.0 with the GTR+F+G4 substitution model. Labels indicate isolate designation, geographic origin (US in black, Ireland in green, Japan in red), and year. Symbols represent pRErm46 plasmid type as indicated in the figure inset (see **Fig. 1** legend). Tree rendered with FigTree (http://tree.bio.ed.ac.uk/software/figtree). (**B**) Accessory genome of MDR-RE 2287 isolates. Gene presence/absence was analysed using Roary v3.12.0 (https://www.sanger.ac.uk/tool/roary/) and visualised with Phandango (https://jameshadfield.github.io/phandango/#/). The two Japanese isolates share similar accessory gene profiles, as do the first two Irish isolates; the third Irish isolate shows a distinct pattern, suggesting a separate introduction event. Eight additional MDR-RE 2287 isolates were included to illustrate representative variation in accessory genome content, along with the prototype MDR-RE 2287 isolate PAM 2287 (chromosome only). The latter serves as control for plasmid-specific accessory gene assignment (labeled “Plasmids”, which includes pRErm46 and the equine-type virulence plasmid, pVAPA). Similarity between isolates was calculated as 2×N_shared_/(N_iso1_+N_iso2_), where N_shared_ is the number of shared accessory genes and N_iso1_/N_iso2_ are the total amount of accessory genes in each isolate, respectively.

The Japanese isolates are located in a distinct sub-branch of the MDR-RE 2287 clonal complex phylogeny, along with the three Irish strains as well as US isolates from Kentucky and New York (**Fig. 2A**). This sub-clonal population shares a characteristic deletion in pRErm46, resulting in loss of the C1I carrying the *aadA9* and *sul1* resistance genes plus the adjacent *tetRA*(33) cassette due to recombination between flanking IS*6100* elements –referred to as the ΔClI-*tetRA*(33) pRErm46 variant [7,8,12] (**Fig. 1**). The consistent presence of the Δ ClI-*tetRA*(33) deletion across this sublineage further supports it is a distinct subclone of MDR-RE 2287. A separate US isolate (strain 146, recovered in Kentucky 2017) (**Fig. 1**) also carries a ΔClI-*tetRA*(33) plasmid variant, showing that the deletion can arise independently in the MDR-RE population. Isolates with a ΔClI-*tetRA*(33) pRErm46 lose resistance to streptomycin, spectinomycin, sulfonamides and tetracycline/doxycycline but retain high-level resistance to macrolides and rifampicin [12].

## Importation events underlying emergence

Within the MDR-RE 2287 ΔClI-*tetRA*(33) subclone, the two Japanese isolates, R22K52 and RE183, are most closely related (**Fig. 2A**), differing by only eight core-genome SNPs, suggesting a common source origin. The same applies to the first two Irish isolates (2016 and 2021), which also differ by just seven SNPs (**Fig. 2A**). In contrast, the average SNP difference across the MDR-RE 2287 clonal complex is 15.24±6.08 SNPs, rising to 35 SNPs between more distantly related isolates. A gene presence-absence analysis of the accessory genome (typically representing ≈7% of an MDR-RE 2287 isolate’s gene content) showed highly similar profiles for the two Japanese isolates and for the two early Irish isolates (**Fig. 2B**), but differences between the two pairs, consistent with their different positioning in the ΔClI-*tetRA*(33) sub-branch of the MDR-RE 2287 clonal complex. These data suggest that the emergence of MDR-RE 2287 in Japan and Ireland each resulted from single importation events of members of the sub-clonal lineage carrying the ΔClI-*tetRA*(33) pRErm46.

In contrast, the 2023 Irish isolate (PAM 2767) differs from the earlier two and clusters more closely with the Japanese and US isolates within the ΔClI-*tetRA*(33) subclone (**Fig. 2A**). Although the limited number of SNP differences constrains resolution, the accessory genome gene content profile further substantiates that the 2023 Irish isolate is distinct from those recovered in 2016 and 2021 (**Fig. 2B**). This suggests that the 2023 isolate likely represents a second, independent importation of a member of the ΔClI-*tetRA*(33) subclonal population into Ireland. The Irish and Japanese strains likely originated in the US, where Δ ClI-*tetRA*(33) MDR-RE 2287 variants have been recovered as early as 2010 (**Fig. 2A**) and are becoming increasingly common (ref. [12] and unpublished data).

## Detection of MDR-RE’s *erm*(46)-carrying transposon in China

As an exploratory approach to assess MDR-RE dissemination, we performed a GenBank search using the MDR-RE-specific 6.9-kb transposon Tn*RErm46*. This identified a hit in *R. equi* strain WH99 (accession ASM3032414v1), isolated in 2015 from soil in Anhui, Eastern China, and studied for its phenazine-1-carboxylic acid (PCA) degradation ability [18,19]. WH99 carries a Tn*RErm46* variant on one of its two plasmids (p1-WH99, NZ_CP121764.1 positions 148,543– 155,119). Unlike the fully conserved Tn*RErm46* sequences in MDR-RE isolates (including non-2287 spillovers), the WH99 copy shows minor divergence (94% identity), including a translocation of the 5’ end to the middle of the transposon sequence (**Fig. 3A**). It is unclear whether this reflects a genuine structural difference or an assembly artefact.

**Fig 3.**
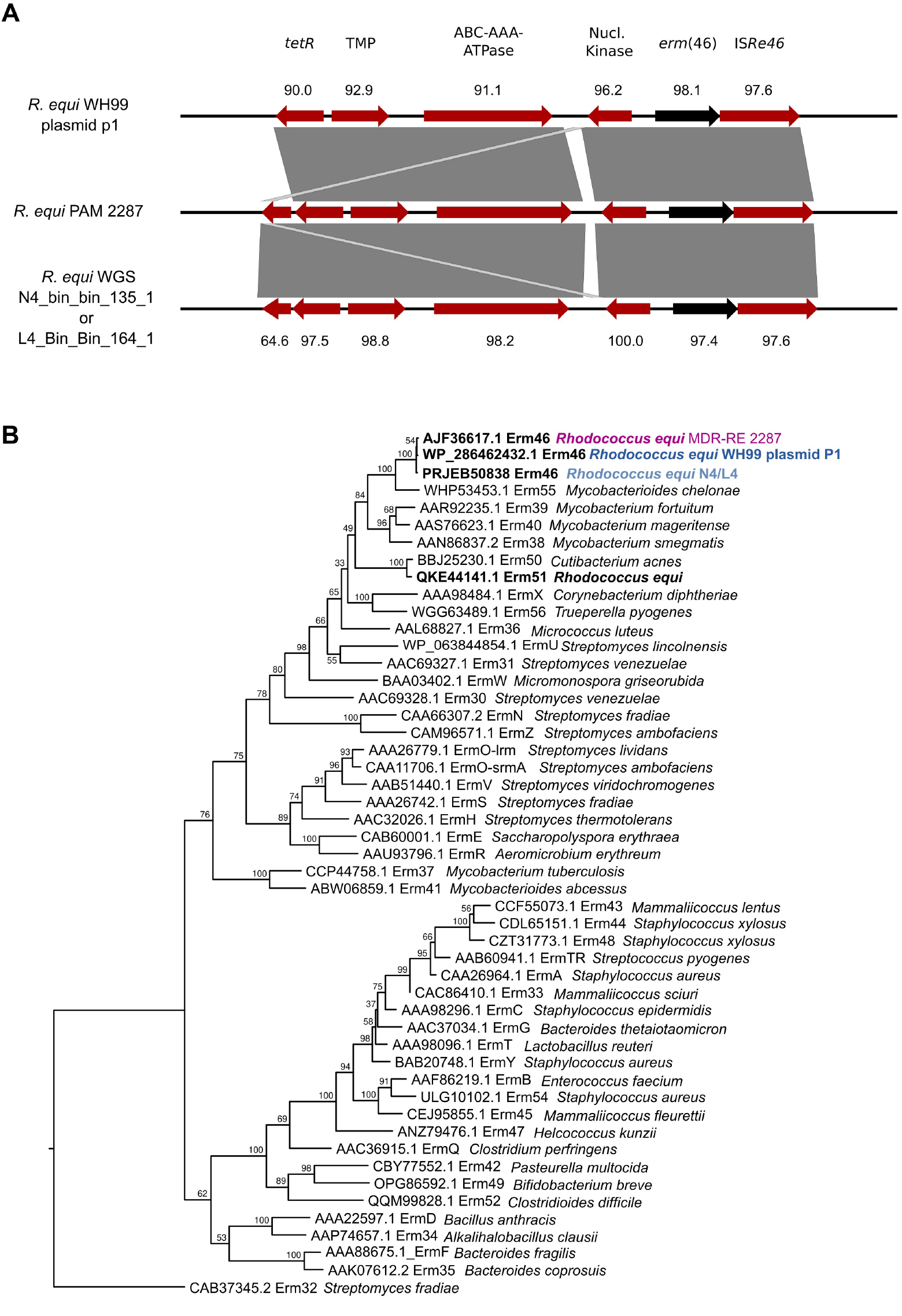
Identification of the MDR-RE transposon Tn*RErm46* in China. (**A**) TblastN alignment of Tn*RErm46* from prototype MDR-RE 2287 isolate PAM 2287 (middle) with Tn*RErm46* sequences from the RefSeq genome assembly of *R. equi* isolate WH99 (top) and the WGS metagenomic assemblies from samples N4_Bin_Bin_135_1 and L4_Bin_Bin_164_1 (bottom). Tn*RErm46* transposon genes are shown in red, except for the macrolide-resistance gene *erm*(46), shown in black. Numbers indicate per cent amino acid identity. (**B**) ML phylogenetic tree based on the amino acid sequences of known Erm 23S methyltransferases, as listed in the MLSB web resource (http://faculty.washington.edu/marilynr/ermweb1.pdf). Protein sequences were aligned using the MAFFT algorithm (G-INS-I, 16 iterations), and phylogeny was inferred with IQ-TREE2 v.2.4.0 using LG substitution model. Ultrafast bootstrap values ≥70% (1,000 replicates) are indicated. The reference Erm46 protein from the *R. equi* MDR-RE 2287 is highlighted in purple, those encoded in the *R. equi* WH99 plasmid P1 or WGS metagenomic assemblies from samples N4_Bin_Bin_135_1 and L4_Bin_Bin_164_1 (both have identical protein sequence) are highlighted in blue and pale blue, respectively. The Erm51 enzyme, another 23S methyltransferase associated with macrolide resistance in *R. equi* [20], is highlighted in bold.

An Erm protein phylogeny confirms that the MDR-RE *erm*(46) product and the WH99 homolog are closely related allelic variants. Both cluster together within the same evolutionary branch alongside mycobacterial Erm proteins (**Fig. 3B**). This is in contrast to the more divergent product of the *erm*(51) gene (47.7% identity) found in a population of *R. equi* soil isolates in the US not linked to clinical disease in horses [20]. *erm*(51) is carried on a distinct transposon mobilized by an IS*605* transposase [20], and its product clusters away from Erm46, near Erm50 from *Cutibacterium* (*Propionibacterium*) *acnes* (**Fig. 3B**). This suggests that the Chinese isolate WH99 carries a slightly diverged version of MDR-RE’s macrolide resistance determinant, potentially shaped by natural genetic drift in an environment lacking horse environment-associated selective pressure.

The detection of the Tn*RErm46* transposon in the Chinese soil isolate WH99 is particularly noteworthy in light of a reported outbreak of foal pneumonia in Beijing in 2013 caused by macrolide/rifampicin-resistant *R. equi* [21]. The causative strain showed MICs ≥25 μg/ml for azithromycin, clarithromycin, and erythromycin, and ≥128 μg/ml for rifampicin [21], matching the MDR-RE phenotype [5,6,12], although it carried a different *rpoB* mutation (H526D) [21]. Transfer of the Tn*RErm46*-harbouring pRErm46 plasmid to diverse *R. equi* backgrounds has been documented, linked with different *rpoB* mutations (e.g. S531L, S531Y) [7,12] likely selected *de novo* by the macrolide-rifampicin combination therapy. Unfortunately, isolates from the Beijing outbreak were not made available for verification, and the paper was retracted by the publisher [22].

Additional evidence supporting the circulation of MDR-RE –or its resistance determinants– in China came from searches of metagenomic datasets. Marine coastal sediments collected in 2019 in Weihai, Eastern China (BioProject PRJEB50838; samples N4_Bin_Bin_135_1 and L4_Bin_Bin_164_1), yielded sequences (CALJQK010000238.1, CALDUI010000034.1) with ≥98% identity and coverage to MDR-RE Tn*RErm46* (**Fig. 3A**). The protein phylogeny in **Fig. 3B** shows that the encoded Erm products cluster closely with those from WH99 and the 2287 clonal isolates, confirming they are allelic variants of MDR-RE’s Erm46.

Apart from those in verified MDR-RE isolates, as well as the Chinese soil isolate WH99 and Weihai metagenomes, no other Tn*RErm46* sequences were detected in the databases at the time of writing.

## Conclusions

The epidemiology of MDR-RE 2287 parallels that of human pathogenic MDR clones. These typically emerge in health-care settings under strong antibiotic pressure, a context comparable to that of horse farms applying systematic *R. equi* antimicrobial prophylaxis. Following emergence and an initial phase of local spread (in the US for MDR-RE), MDR clones spread internationally [23]. Although at a slower pace due to the limited nature of horse movements compared to human travel, the detection of the same subclonal MDR-RE 2287 variant on three continents underscores the risk of *R. equi* becoming globally established. Coordinated international animal and public health surveillance is essential to mitigate this emerging zoonotic AMR threat. A key priority is the molecular screening of horses for MDR-RE carriage before international transport.

## Acknowledgements

We thank our network of international collaborators for their participation in macrolide- and rifampin-resistant *R. equi* monitoring.

Research on multidrug-resistant *R. equi* is supported by the Horserace betting Levy Board (HBLB grant no. vet/prj/814).

